# Long-tailed macaques extract statistical information from repeated types of events to make rational decisions under uncertainty

**DOI:** 10.1101/574475

**Authors:** Sarah Placì, Marie Padberg, Hannes Rakoczy, Julia Fischer

**Author notes:** Corresponding Author: Sarah Placì. Equal contribution (Hannes Rakoczy, Julia Fischer).

## Abstract

Human children and apes are proficient intuitive statisticians when making predictions from populations of objects to randomly drawn samples, whereas monkeys seem not to be. Statistical reasoning can also be investigated in tasks in which the probabilities of different possibilities must be inferred from relative frequencies of events, but little is known about the performance of nonhuman primates in such tasks. In the current study, we investigated whether long-tailed macaques extract statistical information from repeated types of events to make predictions under uncertainty. In each experiment, monkeys first experienced the probability of rewards associated with different factors separately. In a subsequent test trial, monkeys could then choose between the different factors presented simultaneously. In Experiment 1, we tested whether long-tailed macaques relied on probabilities and not on a comparison of absolute quantities to make predictions. In Experiment 2 and 3 we varied the nature of the predictive factors and the complexity of the covariation structure between rewards and factors. Results indicate that long-tailed macaques extract statistical information from repeated types of events to make predictions and rational decisions under uncertainty, in more or less complex scenarios. These findings suggest that, given the right input information, monkeys are intuitive statisticians.

## Introduction

Most individuals live in an uncertain world, characterized by statistical regularities. A key challenge for individuals is to extract these regularities in order to make predictions under uncertainty and adjust their decisions accordingly. Such regularities encompass the non-social as well as the social domain. For instance, during certain times of the year, specific trees are more likely to bear fruits than during other times, while some group members may be more willing to cooperate than others. In addition, different factors may covary conditionally. Your friend may be more willing to lend you money at the beginning of the month than at the end of it, when her budget is low. Trees are more likely to bear fruits not only during the summer, but also when there was sufficient sunshine and rain.

Inferential statistics entails the use of relative frequencies of events, and their covariation patterns with different factors or variables, to derive generalizations about the predictive power of these factors and variables, and to make predictions about future events. This requires the ability to represent factors and variables and to form hypotheses about their relationships with the observed outcomes (Tenenbaum, Kemp, Griffiths & Goodman, 2011). Importantly, statistical inferences rely on calculations describing probabilistic relationships between populations and samples randomly obtained from these populations. This allows to differentiate between patterns of outcomes that are probably the result of chance, and patterns that are probably not and that can therefore be attributed with different confidence levels to hypothesised factors or variables. Inferential statistics allows to strengthen, revise or discard these hypotheses, which in turn increases knowledge about the structure of one’s environment and improves the precision of predictions (Jaynes, 2003).

In the past decades, an increasing body of research has been occupied with exploring the origins of statistical reasoning, and in particular, of intuitions about the probabilistic relationships between populations and randomly drawn samples. Several studies showed that human infants have such intuitions, and use them to make predictions under uncertainty and adjust their decisions accordingly. For example, when observing different types of objects moving in a container, 12-month-olds expect objects of the majority type to randomly exit the container, or objects closer to the exit (Téglás, Girotto, Gonzalez & Bonatti, 2007; Téglás et al., 2011; Téglás, Ibanez-Lillo, Costa & Bonatti, 2015). Similarly, 8- and 12-month-olds use information such as the relative frequencies of static objects present in a container (Xu & Garcia 2008), as well as information about sampling conditions (random or biased) (Xu & Denison, 2009), to predict which type of object is going to be sampled from the recipient, and to adjust their decisions accordingly (Denison & Xu, 2014).

Infants also seem to use probabilistic intuitions to make predictions and generalizations from samples to populations. For example, 6- and 8-month-olds integrate prior knowledge about the possible distribution of objects in different populations, and evidence produced by samples obtained from a hidden population, to predict the content of that population (Xu & Garcia, 2008, Denison, Trikatum & Xu, 2014). Note, however, that controls to rule out the possibility that children relied on quantity heuristics rather than probabilities were not conducted in these studies. Fifteen-month-olds rely on cues of random or biased sampling (based on the probability of obtaining the observed sample given a random or a biased sampling, or based on the behaviour of the agent performing the sampling), in order to flexibly generalize the property of the sampled objects to only part of the population, or to the entire population (Gweon, Tenenbaum & Schulz, 2010). Taken together, these findings suggest that some intuitive form of statistical reasoning evolved independently from human culture as a powerful induction mechanism allowing humans, from very early on, to infer meaning from the probabilistic contingencies structuring their physical and social world.

To explore whether intuitive statistics is evolutionary ancient and shared with our closest relatives, apes (Eckert, Call, Hermes, Herrmann & Rakoczy, 2018; Eckert, Rakoczy, Call, Herrmann & Hanus, 2018; Rakoczy et al., 2014), as well as New and Old World monkeys (Tecwyn, Denison, Messer & Buchsbaum, 2017; Placì, Eckert, Rakoczy & Fischer, 2018, respectively) were tested for their ability to predict the outcomes of sampling events out of populations of objects, based on a paradigm previously developed for human infants (Denison & Xu, 2014). While results from the ape studies indicate that they, similar to human infants, possess this ability, results from the monkey studies were more ambiguous. The capuchin monkeys did not perform well in a control condition necessary to rule out the use of a quantity heuristic in their reasoning process, and among the group of long-tailed macaques, only one individual performed well in most experiments, whereas the rest of the group seemed to rely on quantity heuristics. Similar to human infants, apes consider sampling conditions (random or biased) when making inferences from populations to samples (Eckert et al., 2018a). Apes were also compared to human infants in their ability to reason from samples to populations of objects (Eckert, Rakoczy & Call, 2017). The results of this study indicate that the apes based their choices on the absolute number of objects present in samples, rather than on the probability that a given population produced a given sample. These findings suggest that apes possess similar statistical reasoning abilities to human children, but that monkeys might rely on quantity heuristics to make predictions.

However, the ability to rely on statistical information to make predictions under uncertainty need not necessarily be investigated with paradigms akin to games of chance, in which objects are randomly sampled out of a container. Different studies investigated children’s ability to extract statistical information from repeated types of events to estimate the likelihood with which different factors produce different frequencies of outcomes. Findings suggest that human children are also proficient intuitive statisticians in such tasks. For example, 24-month-olds rely on the frequencies of repeated actions and their outcomes (e.g., putting an object on a device – the device is activated or not), and on how different frequencies of outcomes covary with different factors (e.g., different objects put on the device), to infer which factor is more likely to bring about the outcome (Waismeyer, Meltzoff & Gopnik, 2015). Moreover, when an outcome is conditional on more than one factor (e.g., music being produced by a toy manipulated by a social agent), 16-month-olds can infer which factor failed to cause the desired outcome (e.g., music) by relying on the covariation structure between outcomes and factors (Gweon & Schulz, 2011). More particularly, when the production/non-production of music covaried with different toys, children inferred that the toys were responsible for not producing music, and when it covaried with different social agents, children inferred that the social agents were responsible for the non-production of music. Finally, 2.5-years-olds relied on statistical dependencies of repeated types of events and their outcomes (e.g., relative frequencies at which two different objects activate a mechanical device together or separately), in order to predict which object is more likely to activate a device (Gopnik, Sobel, Schulz & Glymour, 2001; Völter, Sentís & Call, 2016).

Apes and capuchin monkeys were also tested in paradigms in which they had to make predictions based on statistical information sampled from repeated events (Völter et al. 2016, Edwards et al., 2014). Findings suggest that they relied on the relative frequencies at which two different objects activated a mechanical device together or separately, to predict which object was more likely to activate a device on its own. However, all these studies did not systematically test whether subjects relied on statistical information rather than on quantity heuristics.

In summary, these findings suggest that some aspects of statistical reasoning are evolutionary ancient. While apes share some (but not all) of the abilities of human children, monkeys are faring less well than apes. It is however not fully clear, whether monkeys would be able to process statistical information extracted from repeated types of events rather than on quantity heuristics, in order to make predictions under uncertainty. In other words, the presentation format may matter in the application of intuitive statistical abilities. The current study aimed to shed more light on this question. More specifically, we aimed to investigate whether a group of long-tailed macaques (*Macaca fascicularis*), that previously failed in making predictions from populations of objects to samples (see Placì et al. 2018), would be able to extract statistical information from a succession of repeated actions, to make predictions about new events and adjust their decisions accordingly. We also aimed to investigate the boundaries of such ability, by varying the nature and the number of factors potentially producing the outcomes.

The study contained three main experiments. In Experiment 1, monkeys had the choice between two objects that were associated with different probabilities to yield a reward. In a sampling phase, monkeys experienced the statistical patterns of rewards associated with both objects separately. In the subsequent test, monkeys could then choose between the two objects presented simultaneously. Different conditions were used to rule out that monkeys relied on quantity heuristics. In Experiment 2, monkeys were presented with two humans operating one box containing grapes. The two persons differed in their propensity to open the box and reward the monkeys with a grape. Again, monkeys experienced the rewarding patterns associated with both humans separately in the sampling phase, before having to make a choice between both humans in the test trial. In Experiment 3, monkeys were presented with two humans operating two boxes. The rewarding pattern were conditional either on the boxes or on the humans depending on the condition. In a first condition, the rewarding patterns covaried with the two boxes in the sampling phase (both humans were better at opening one box than the other), and in the second one, with the two humans (one human was better at opening both boxes). Monkeys could therefore learn in the sampling phase, whether the humans or the boxes mattered for their subsequent choice during the test trial.

### Experiment 1

The aim of this experiment was twofold. We first aimed to assess whether long-tailed macaques extract numerical information from repeated types of events at all, in order to make rational decisions. Second, we aimed to assess whether their decisions are based on probabilities (comparing relative frequencies) or on quantity heuristics (comparing absolute frequencies). Experiment 1 consisted of four test conditions (Exp.1a-d) divided in twelve sessions, in which long-tailed macaques could first learn rewarding frequencies associated with two objects, and then choose between both objects presented simultaneously. Exp. 1a-c differed in respect to the rewarding frequencies associated with both objects (see Table 1). Exp. 1a tested monkeys’ ability to select the best option in an easy scenario, in which the favourable object was probabilistically and absolutely associated to more rewards (favourable object: 10 out of 10 rewards; unfavourable object: 4 out of 10 rewards, see Table 1).

**Table 1.** Rewarding patterns. This table shows for each condition, the number of interactions monkeys had with each option, and the number of associated rewards.

|  | Number of interactions / number of rewards received |  |
| --- | --- | --- |
|  | Favourable option | Unfavourable option |
| Exp. 1a | 10/10 | 4/10 |
| Exp. 1b | 8/10 | 4/10 |
| Exp. 1c | 4/4 | 4/10 |
| Exp. 1d | 4/4 | 4/10 |
| Exp. 2a | 6/6 | 6/12 |
| Exp. 2b | 4/4 | 4/10 |
| Exp. 3a | 10/10 | 4/10 |
| Exp. 3b | 10/10 | 4/10 |

Success in such a scenario could be based on comparing probabilities, on comparing only absolute frequencies, or on comparing certain to uncertain patterns. Exp. 1b aimed to rule out this latter strategy, by associating both objects with uncertain rewarding frequencies (favourable object: 8 out of 10 rewards; unfavourable object: 4 out of 10 rewards). Success in such a scenario could still be based on a comparison of absolute frequencies. Exp. 1c aimed to rule out this strategy, by associating both object with the same absolute number of rewards, but with different probabilities (favourable object: 4 out of 4 rewards; unfavourable object: 4 out of 10 rewards). In Exp. 1a-c, the side on which both objects were presented was kept constant during all twelve session. Exp. 1d was similar to Exp. 1c in terms of the rewarding patterns associated with both options, but not in terms of the side on which they were presented, which was counterbalanced between sessions. The aim of this conditions was to control whether long-tailed macaques really chose between the two objects rather than between the two sides.

## Methods

### Subjects

13 monkeys – aged 1 to 8 years – took part in Exp. 1a but only 12 (female N = 5) completed it (see Table S1 of the supplementary material for details). 10 monkeys (female N = 4) took part and completed Exp. 1b, 11 subjects took part in Exp. 1c but only 10 (female N = 4) completed it and 13 subjects (female N = 7) participated and completed Experiment 1d. The monkeys lived in a large social group of 35 individuals. They were housed at the German Primate Center in Göttingen, Germany, and had access to indoor (49 m^2^) and outdoor areas (173 m^2^), which were equipped with branches, trunks, ropes and other enriching objects. All individuals were already experienced in participating in cognitive experiments and some of them previously took part in experiments requiring them to indicate a choice between two objects via pointing or reaching towards it. All testing was non-invasive, and subjects participated voluntarily. They were not food deprived for testing, and water was always available ad libitum. The monkeys were fed regular monkey chow, fruits and vegetables twice a day.

### Ethics

All experiments were performed under the control of experienced veterinarians to ensure that the studies were in accordance with the NRC Guide for the Care and Use of Laboratory Animals and the European Directive 2010/63/EU on the protection of animals used for scientific purposes. In accordance with the German Animal Welfare Act, the study was approved by the Animal Welfare Officer of the German Primate Center: according to the German Law, the experiments are not invasive and do not require permission by higher authorities (LAVES Document 33.19-42502).

### Experimental Setup

Monkeys were tested in a testing cage (2.60 m × 2.25 m × 1.25 m; height × width × depth) that was adjacent to their indoor enclosure, and could be subdivided into six experimental compartments. Subjects were tested individually in one compartment (1.05 m × 1.10 m; height × length) to which an attachable cage (73 cm × 53 cm × 35 cm; height × width × depth) was fixed, allowing subjects to have a better access to the experiment. The cage was built in metallic mesh, except for the front part that separated the monkey and the experimenter, which consisted of a removable Plexiglas pane (27 cm × 34 cm; height × length). The pane had two small holes (ᴓ 3.5 cm; distance between holes 27 cm; see Fig. 1) through which subjects could insert their arm to indicate a choice. A wheeled table (85 cm × 80 cm × 50 cm; height × width × depth) was set in front of the cage, on which different objects could be displayed. The experimenter (or the two humans) stood behind the table.

**Fig. 1.**
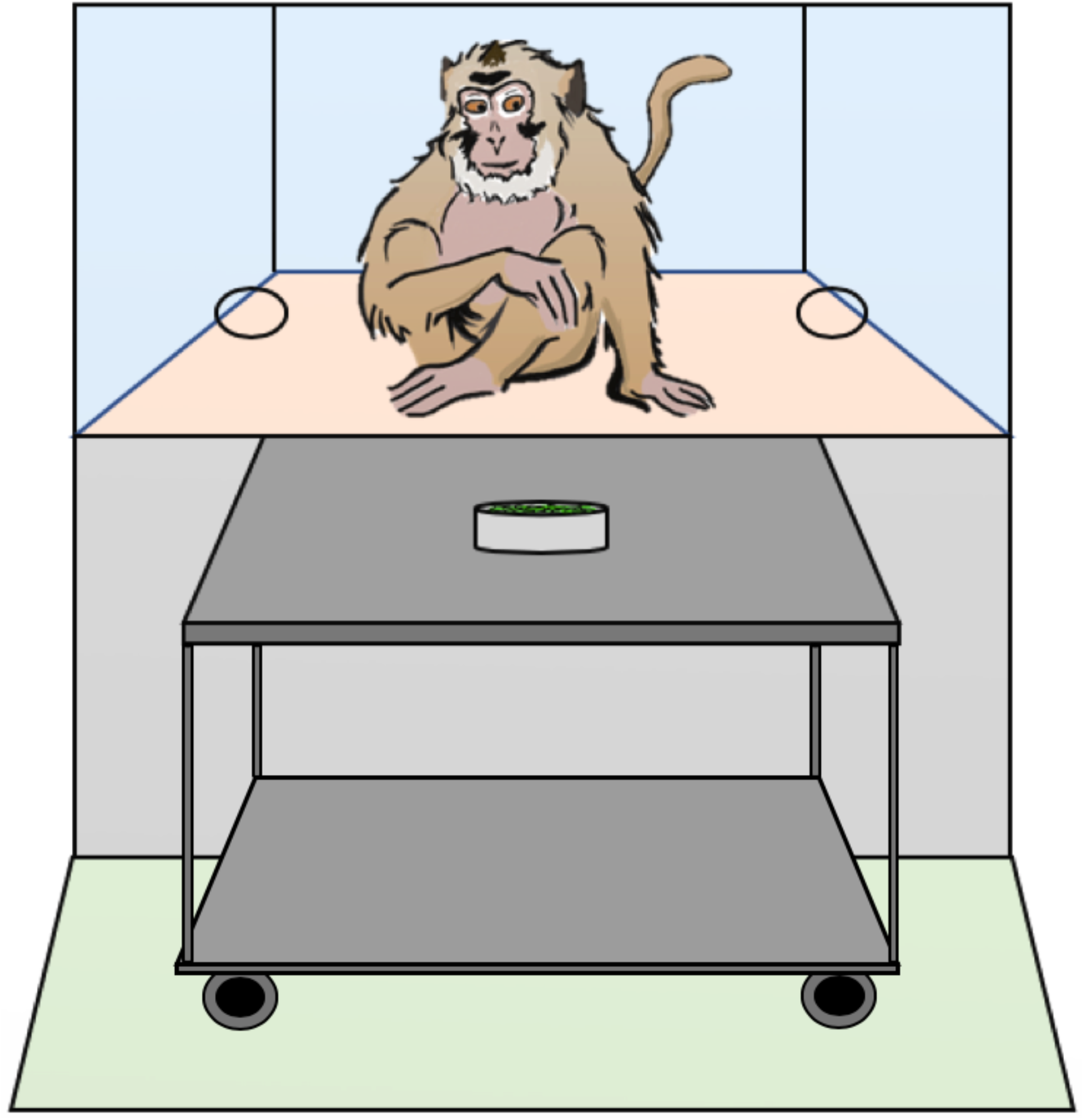
Experimental setup. Subjects stood in an extension of the testing cage. A Plexiglas panel with two holes allowed them to touch the objects or the hands that were presented to them. A wheeled table stood in front of the cage, on which the testing material was placed, and behind which stood the experimenter (Exp. 1) or the different social agents (Exp. 2 and 3).

### Study procedure

Each test condition consisted of 12 sessions. Each session was subdivided into a sampling phase, and a test trial (see Fig. 2). In all conditions, the options presented to subjects were different coloured climbing carabiners. Each of the 12 sessions unfolded as follow: the experimenter would approach the first carabiner towards one of the two holes in the Plexiglas, wait for the monkey to touch it, and then reward the monkey with a grape or not, from a bowl standing in the middle of the table (see Figure 1). This procedure was repeated four or ten times depending on the option and the condition (see Table 1), with 20 seconds intervals between the moment the monkey touched the carabiner and the next presentation. The same procedure was repeated for the second carabiner (sampling phase). After the last presentation, the experimenter would wait 30 seconds and then present both carabiners simultaneously (test trial). Rewarding frequencies associated with both objects in the test trial were meant to reflect those in the sampling phase, but as there was only one trial per session, they were spread across the twelve test trials of the twelve sessions. All non-rewarded events were pseudo randomly distributed among all presentations. All subjects started with Exp. 1a, half of them went on with Exp. 1b and then Exp. 1c, and half with Exp. 1c and then Exp.1b. All subjects completed Exp. 1d last. In all experiments, all sessions were separated by either half a day or more. In each condition, the option presented first was counterbalanced across trials, and the colour of option A was counterbalanced between subjects. Different combinations of coloured carabiners were used in the different conditions.

**Fig. 2.**
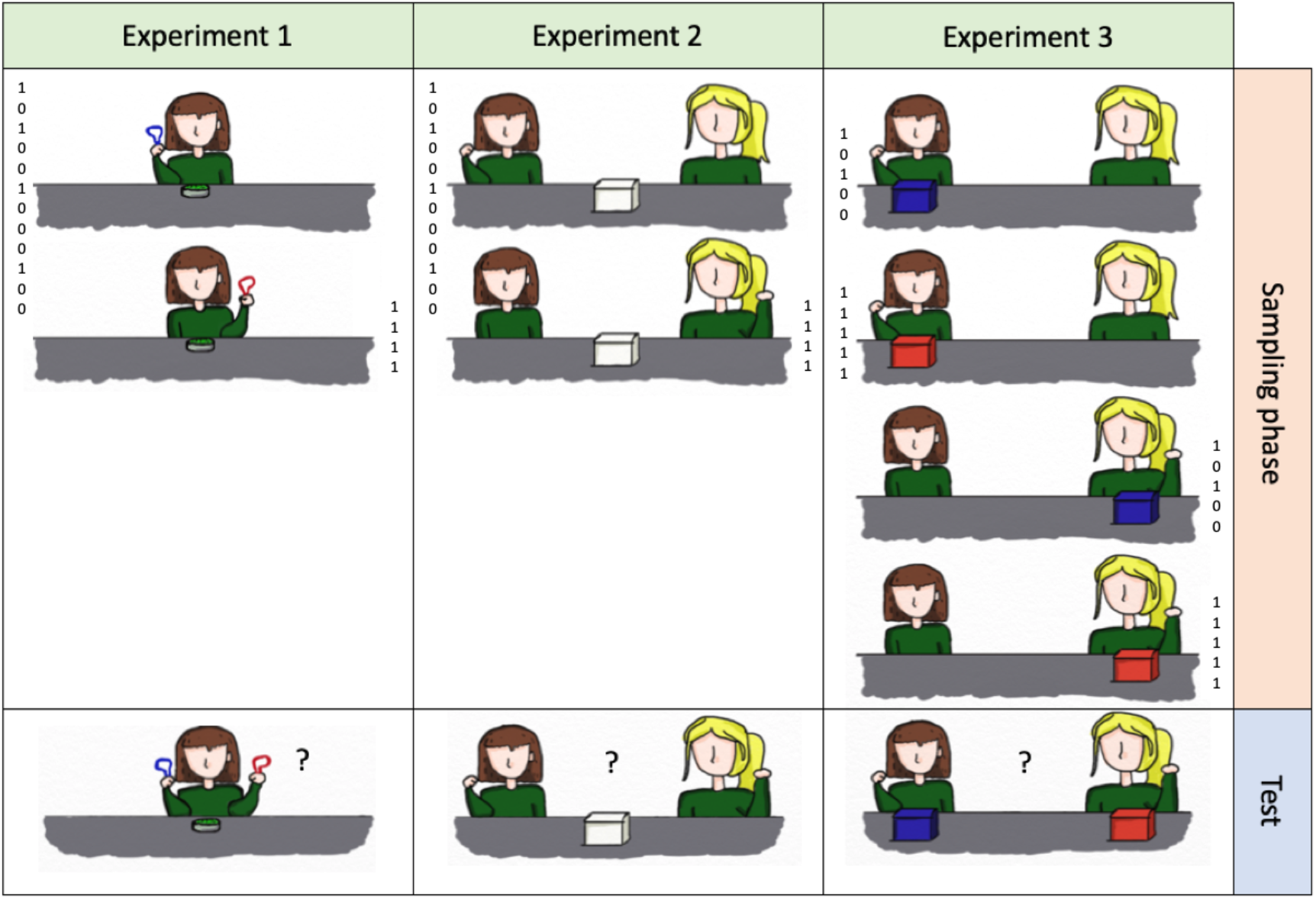
Example of the procedure of one session for each experiment. In Experiment 1, the experimenter would approach a first object to a hole in the Plexiglas and wait for the monkey to touch it, and then reward the monkey or not. This would be repeated a certain number of times with the first object, and then with the second (sampling phase). The numbers (1 or 0) represent an example of rewarding pattern of Exp. 1c and 1d. In the test trial, both objects would be presented simultaneously, and the monkey would have to touch one in order to indicate her choice. In Experiment 2 and 3, first one human, then the other would repeatedly present her fist to the monkey and then try to open the box. If she succeeded, she would grab a grape in the box and hand it to the monkey. The numbers (1 or 0) are an example of the rewarding pattern in Exp. 2b and Exp. 3b, and indicate the success of the human and the resulting reward.

### Coding and analysis

Every session was video recorded. The experimenter coded monkeys’ choices live. Whenever monkeys chose the favourable option, we scored it as a success, while we scored it as a failure when they chose the unfavourable option. A second observer coded 25% of the sessions of the first three conditions, using the video recordings. Agreement between the experimenter and the second coder was perfect for all experiments (100% of agreement). To test whether monkeys’ mean group performance for each experiment was different from chance, we computed one sample one-tailed t-tests. In addition, to check whether monkeys’ performance changed over the course of sessions, we computed the correlation coefficient between group performance and trial number separately for all conditions. All analyses were performed using R (R core Team 2017).

## Results and discussion

On average, on a total of 12 test trials, subjects selected the favourable option in 10.7 trials in Exp. 1a (88.9%; *SD* = 0.78 trials), 9.6 trials in Exp. 1b (80%; *SD* = 1.58 trials), in 8.4 in Exp. 1c (70%; *SD* = 2.91 trials), and in 8.3 trials in Exp. 1d (69.2%; *SD* = 1.65 trials), significantly more than expected by chance in all conditions (Exp. 1a: *t*(11) = 20.77, *p* < .001, *d* = 5.99; Exp. 1b: *t*(9) = 7.22, *p* < .001, *d* = 2.28; Exp. 1c: *t*(9) = 2.61, *p* = .01, *d* = 0.82; Exp. 1d: *t*(12) = 5.04, *p* < .001, *d* = 1.40, see Fig. 3, for individual performances, see Table S1 for the supplementary material). In addition, in all conditions, there was a significant correlation between group performance and trial number (Exp. 1a: *r* = 0.77, *p* < 0.01; Exp. 1b: *r* = 0.86, *p* < .001; Exp. 1c: *r* = 0.97, *p* < .0001; Exp. 1d: *r* = 0.7, *p* = .01, see Fig. 4). Long-tailed macaques reached ceiling in the last trials of all conditions except in Exp. 1d. These results suggest that long-tailed macaques as a group extracted statistical information provided in the sampling phase, to estimate the probability with which interacting with each object would yield a reward, and adjusted their decisions accordingly in the test trials. Monkeys good performance in Exp. 1d further indicates that they considered the objects to be predictive of rewards, and not only the side on which they were presented. Interestingly, even if monkeys performed above chance in all conditions, the rewarding patterns associated to both options – and therefore the condition – seemed to have an effect on their performance. We ran a post-hoc Generalized Linear Mixed Model (GLMM; Baayen, 2008), with monkeys’ choices in Exp. 1a, 1b and 1c as dependant variable, condition, trial number and the order in which each individual underwent the different conditions as fixed factors, and individual ID as random factor, to further test if the condition had an effect on monkeys’ performance, and to assess whether there were differences between conditions. The GLMM revealed that condition had a main effect on subject’s performance (*X*^2^ = 14.457, *df* = 2, *p* = 0.001; see Table S2 of supplementary materials for more details), as well as the trial number. A post-hoc investigation showed that monkeys performed significantly better in Exp. 1a (favourable object: 10/10 rewards; unfavourable object: 4/10 rewards) compared to Exp. 1c (favourable object: 4/4 rewards; unfavourable object: 4/10 rewards; estimate ± *CI* = −2.31 ± 0.69, p < .01, see supplementary materials for more details), and in Exp. 1b (favourable object: 8/10 rewards; unfavourable object: 4/10 rewards) compared to Exp. 1c (estimate ± *CI* = 0.98 ± 0.37, p = .02). These results suggest that long-tailed macaques considered both the relative frequency of rewards associated with both objects, and the number of presentations of each object (and therefore the sample size) in the sampling phase, to make predictions and direct their choices in the test trial. The fact that their performance in choosing the favourable option improved across sessions in every condition further suggests that the confidence of their predictions increased with an increased sample size, as with each new session, they underwent a new sampling phase. An alternative explanation could, however, also account for these results, namely that long-tailed macaques relied on a strategy of avoidance of the object that was associated with the most non-rewarding events. Yet, if this was the case, they should have performed at similar levels in Exp. 1a (10/10 vs. 4/10) and Exp. 1c (4/4 and 4/10), which they did not.

**Fig. 3.**
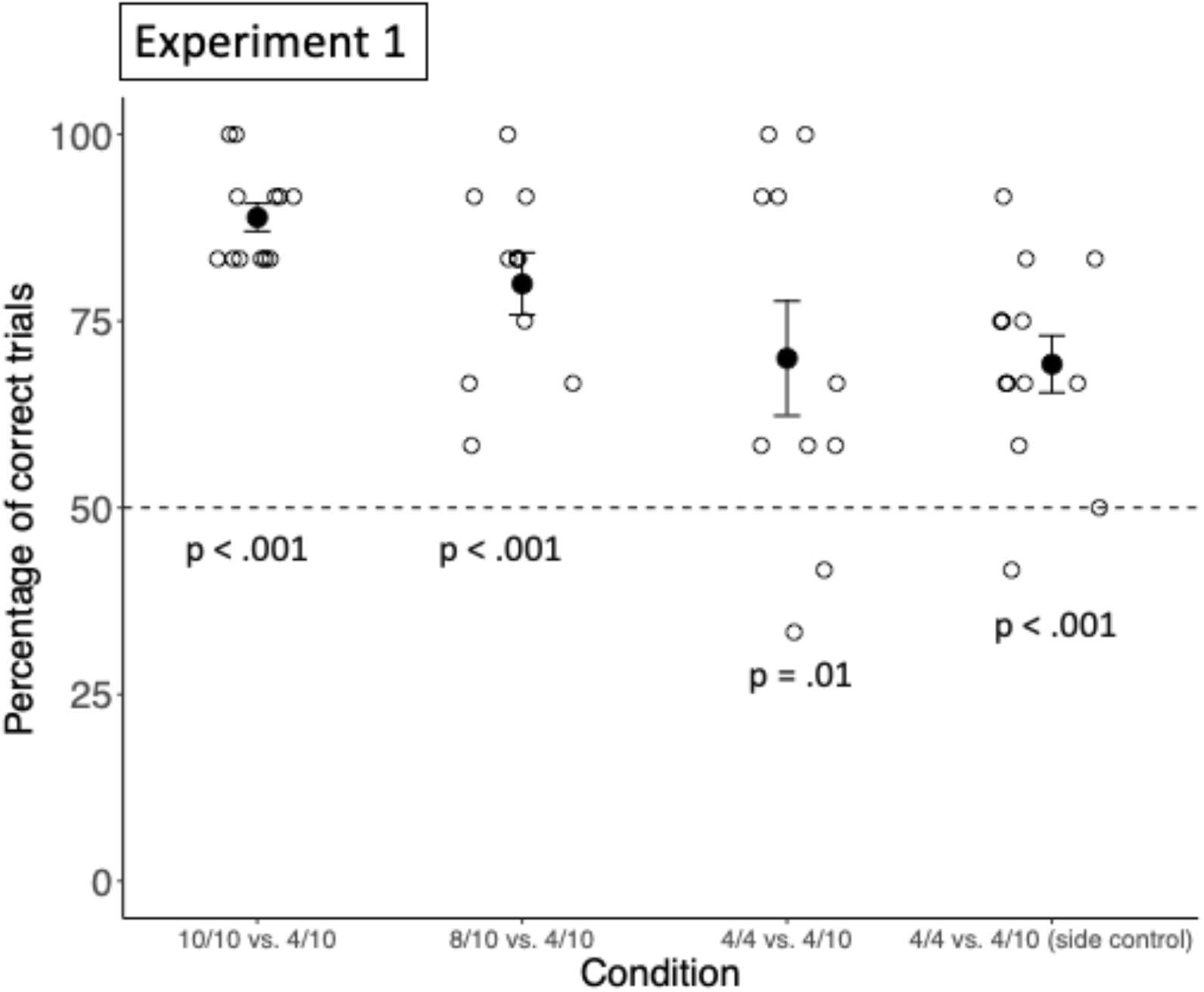
Group performance in Experiment 1. This figure shows the group mean percentage of choosing the option leading to more probable rewards, in each condition. Filled circles represent the group means, and the black back bars represent the standard errors. The empty circles represent individual performances.

**Fig. 4.**
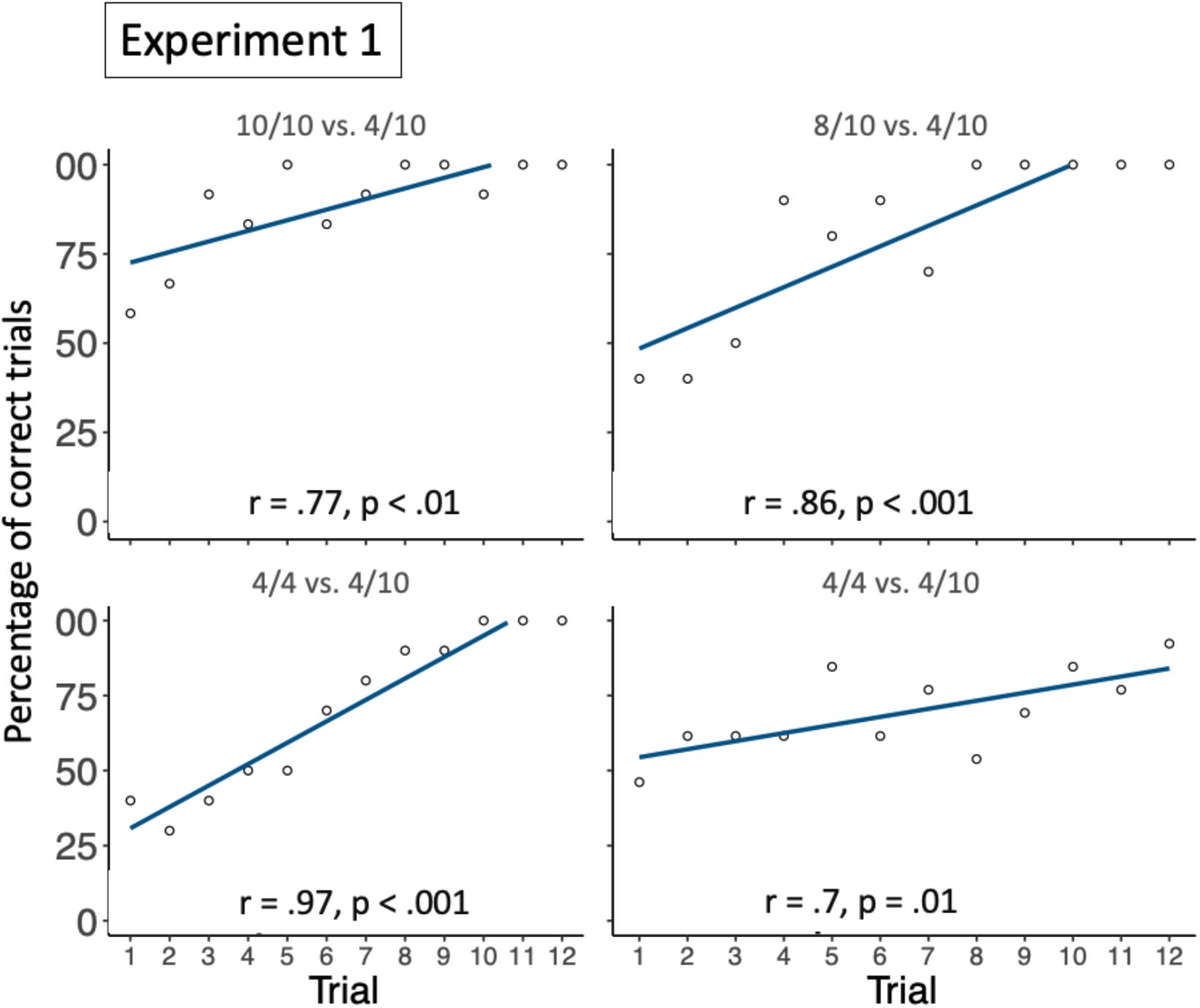
Group performance across trials in Experiment 1. This figure shows the correlation between group performance and trial number for each condition. The dark lines are regression lines and the light lines show the closest fit to the data points.

### Experiment 2

Every day uncertainty also pertains to what others will do, especially for highly sociable species. In Experiment 2, we aimed to further investigate the boundaries of long-tailed macaques statistical reasoning abilities, by exploring whether they can also extract statistical information from and make predictions about more complex scenarios involving social agents performing an action. In this experiment, rewarding events were caused by two humans trying to open a box containing grapes. If they succeeded in doing so, they would grab a grape and give it to the monkey, if they did not succeed, they would stop trying and the monkey would receive no reward.

Experiment 2 consisted of two conditions (Exp. 2a and Exp. 2b) subdivided into twelve sessions. Both conditions only differed in terms of the rewarding patterns associated with both agents. We started by running Exp. 2a and decided to test a rewarding pattern similar to Exp. 1c and 1d, but with more presentations in the sampling phase (favourable agent: succeeded to open the box 6 times out of 6 attempts; unfavourable agents succeeded 6 times out of 12 attempts). We then run Exp. 2b with similar rewarding patterns as in Exp. 1c and 1d, to have a better comparison (favourable agent: succeeded 4 times out of 4 attempts; unfavourable agents succeeded 4 times out of 10 attempts).

## Methods

### Subjects

12 subjects (female N = 6) participated and completed Exp. 2a and 15 subjects participated in Exp. 2b but only 14 (female N = 8) completed it (see Table 1 for more details). Monkeys were part of the same social group and the same housing conditions as in Experiment 1.

### Ethics

See Experiment 1.

### Study procedure

Each of the 12 sessions unfolded as follow: a white box (10 cm × 10 cm × 10 cm) containing grapes and closed by a sliding lid was set in the middle of the table. Behind the table stood two humans. Following the instructions of the experimenter, one human started the session by moving her closed fist towards the hole in the Plexiglas behind which she stood, and waited for the monkey to touch it. Once the monkey touched her, she tried to open the white box with the help of a little L-shaped tool (Allen key). If she succeeded (successful trials were communicated by the experimenter), she would grab a grape from within the box and hand it to the monkey as reward. If she did not succeed, she would try a bit longer and then stop, so as to keep successful and unsuccessful trials of equal length. This was repeated a certain number of times with twenty seconds intervals between the moment the monkey touched the fist and the next presentation, after what the second person started the same procedure (sampling phase). Thirty seconds after the completion of the sampling phase, on the signal of the experimenter, both humans moved their fists simultaneously to their corresponding hole in the Plexiglas, and waited for the monkey to touch one of them (test trial, see Fig.2). All non-rewarded trials were pseudo randomly distributed among all presentations. All sessions were separated by a day or more. In both conditions, the side on which agent A stood was counterbalanced across sessions. In each condition, the person who started the session was counterbalanced across sessions, and the identity of agent A was counterbalanced between subjects. The two social agents were changed between conditions.

### Coding and analysis

See Experiment 1. In this experiment, we refrained from further observer reliability assessments, as agreement was perfect in Experiment 1.

## Results and discussion

On average, subjects selected the favourable agent in 7.7 trials in Exp. 2a (63.9%; *SD* = 1.50 trials) and in 6.93 trials in Exp. 2b (57.8%; *SD* = 1.83 trials), significantly more than expected by chance (Exp. 2a: *t*(11) = 3.86, *p* < .01, *d* = 1.11; Exp. 2b: *t*(13) = 1.83, *p* = .05, *d* = 0.51, see Fig. 5, for individual performances, see Table S1 for the supplementary material). There was however no significant correlation between group performance and trial number (Exp. 2a: *r*= 0.024, *p*= 0.94; Exp. 2b: *r*= 0.07, *p* = 0.84, see Fig. 5). These results suggest that long-tailed macaques as a group extracted statistical information from more complex events involving social agents to estimate the probability with which interacting with each person would yield a reward, and adjusted their decisions accordingly. However, their lack of learning in Exp. 2b, compared to Exp. 1d, indicates that despite the same rewarding frequencies associated with both options, Experiment 2 was harder than Experiment 1. There are several possible reasons explaining this pattern of behaviour. First, because of the increased complexity of the task (two social agents manipulating tools to open a box containing rewards), monkeys might have needed more than 12 sessions to be confident about which factor lead to more probable rewards and were still considering different possibilities. Second, it could be that monkeys did not recognize that the two humans were the same individuals across all sessions and therefore did only estimate their predictive power within but not between sessions. This, however, is not very likely, at least in Exp. 2a, because the monkeys knew the two humans well. Third, it might be that long-tailed macaques do not represent humans’ propensity to open a box as being stable over time, and therefore treated each session separately. Fourth, because of the complexity of the task, it is possible that this time, monkeys only considered the predictive power of the side of presentation of both humans, and not of the humans themselves. As the side was only predictive within but not between sessions (i.e., the same human would stand on the same side of the table during the sampling phase and the test trial, but the side was counterbalanced across sessions), it could be a reason why their performance was above chance level but did not increase over sessions.

**Fig. 5.**
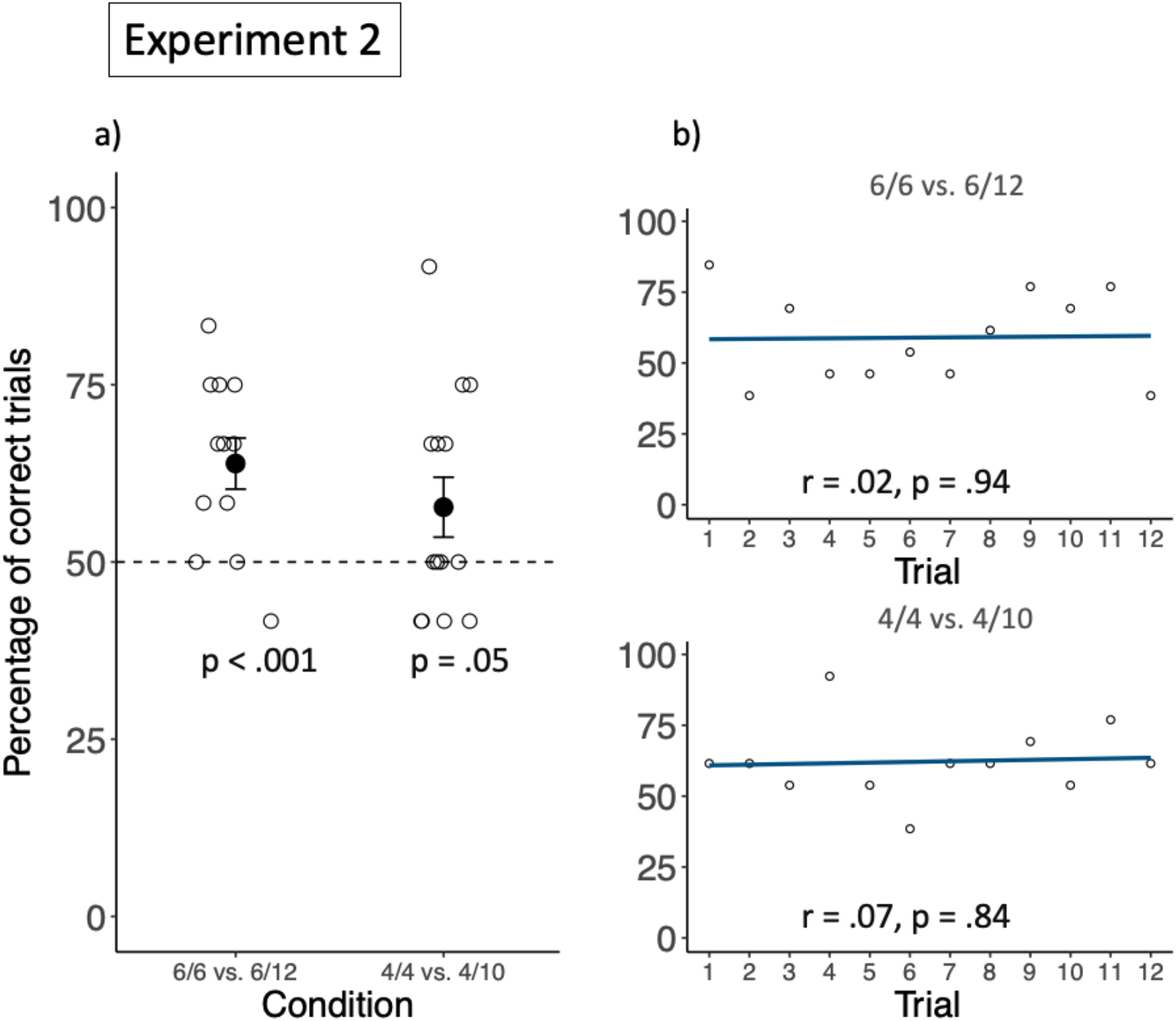
Group performance in Experiment 2. a) shows the group mean percentage of choosing the favourable option, in both conditions. Filled circles represent the group means, and the black back bars represent the standard errors. The empty circles represent individual performances. b) shows the correlation between group performance and trial number for each condition. The dark lines are regression lines and the light lines show the closest fit to the data points.

### Experiment 3

In Experiment 3, we aimed to further investigate the boundaries of the long-tailed macaques’ statistical reasoning abilities in repeated types of events scenarios. More particularly, we aimed to explore whether long-tailed macaques can also extract statistical information from complex scenarios involving both social and non-social factors, and in which the rewarding frequencies are either conditional on the social or on the non-social factors. In real life scenarios, when an agent is good at performing a task, it could be due to intrinsic characteristics of that agent or of the task. By monitoring covariation patterns between different agents and different tasks, one can disentangle between the different possibilities (see Gweon & Schulz, 2011, for an example with human infants). In Experiment 3, rewarding events were also caused by two humans, but this time both humans would try to open two different boxes. Therefore, the different humans and the different boxes could potentially account for the variation in the rewarding patterns. Experiment 3 consisted of two conditions. In Exp. 3a, one human was more likely to open both boxes (she would succeed 5 times out of 5 with both boxes – 10 times out of 10 times in total – compared to the other human who would succeed 2 times out of 5 with both boxes – in total 4 times out of 10), but both boxes would be opened at the same rates. In Exp. 3b, one box was more likely to be opened by both humans (one box would be opened by both humans 5 times out of 5 – 10 times out of 10 in total – whereas the other would only be opened 2 times out of five by both humans – in total 4 times out of 10, see Table 1), but both humans would open boxes at similar rates. In the test trial, one box was put in front of one human, and the other box in front of the other human. If long-tailed macaques were able to pay attention to the covariation pattern between the rewarding frequencies and the different factors (humans or boxes), we predicted that in Exp. 3a, they would base their decisions on the human, irrespective of the box placed in front of her, whereas in Exp. 3b, they would base their decisions on the box, irrespective of the human standing behind it. This experiment additionally allowed us to test more directly whether monkeys’ lack of learning in Experiment 2 was due to their inability to consider social agents as predictive factors across sessions. If this was the case, their performance should increase across sessions in Exp. 3b, in which the boxes were predictive of the different rewarding patterns, but not in Exp. 3a, in which the social agents were predictive.

## Methods

### Subjects

Twelve and eleven subjects participated in and completed Exp. 3a and Exp. 3b respectively. Monkeys were part of the same social group and the same housing conditions as in Experiment 1.

### Ethics

See Experiment 1

### Study procedure

The procedure was the same as in Experiment 2, except that instead of trying to open only one box, each human tried to open two boxes differing in colour (See Fig. 2). The boxes were this time not placed in the middle of the table, but presented one after the other in front of each human, to allow both humans to open both boxes. Each box therefore appeared on both sides of the table during the sampling phase. In the test phase, both humans kept their places, and one box was put in front of each human. Sessions were separated by a day or more. In both conditions, the side on which the boxes were placed during the test trial was counterbalanced between sessions. In both conditions, the side on which person A stood was counterbalanced across sessions. In each condition, the person who started the session was counterbalanced across sessions, and the identity of the favourable agent was counterbalanced between subjects. Trials in which the first box was first opened by both persons, and trials in which one person first tried to open both boxes, were counterbalanced between sessions. All non-rewarded trials were pseudo randomly distributed among all presentations. The social agents were changed between conditions. Half of the subjects started with Exp. 3a and then Exp. 3b, and the other half started with Exp. 3b first. Approximately 20 days elapsed between both conditions.

### Coding and analyses

See Experiment 1. In this experiment, we refrained from further observer reliability assessments, as agreement was perfect in Experiment 1.

## Results and discussion

On average, subjects selected person A in 7.1 trials (59%; *SD* = 0.58 trials) in Exp. 3a, and box A in 7.36 trials (61.4%; *SD* = 1.00 trials) in Exp. 3b, significantly more than expected by chance (Exp. 3a: *t*(10) = 2.00, *p* = .04, *d* = 0.58; Exp. 2b: *t*(10) = 3.32, *p* < .01, *d* = 1.00, see Fig. 6, for individual performances, see Table S1 for the supplementary material). In addition, in both conditions, there was no significant correlation between group performance and trial number (Exp. 3a: *r* = 0.48, *p* = 0.12; Exp. 3b: *r* = 0.13, *p* = 0.68, see Fig. 6). These findings suggest that long-tailed macaques as a group relied on more complex covariation structures, to estimate which factor would more likely lead to a reward. Furthermore, these findings suggest that the difference in monkeys’ performance between Experiment 1 and 2 was not due to their inability to rely on humans as predictors, as their performance was also lower in Exp. 3b, and did not increase across trials. Also, it seems unlikely that their above chance performance, but lack of learning across sessions in Experiment 2 and 3 was due to their reliance on the side of presentation as predictor rather than on the humans or the boxes. If this was the case, long-tailed macaques should have performed at chance level in Exp. 3b, because the side on which the different boxes were presented in the sampling phase was not predictive of their side of presentation in the test phase. Their overall lower performance and lack of learning in Experiment 2 and 3, compared to Experiment 1, might therefore have been due to the increased complexity of the tasks. As there were more potential factors to consider, monkeys might simply have needed more time to learn with confidence which factor was predicting a more likely reward across sessions. Alternatively, monkeys could also have had trouble to represent different abilities to open boxes, and different propensities of boxes to be opened as enduring characteristics of social agents and boxes and might not have extended their learning from one session to the other.

**Fig. 6.**
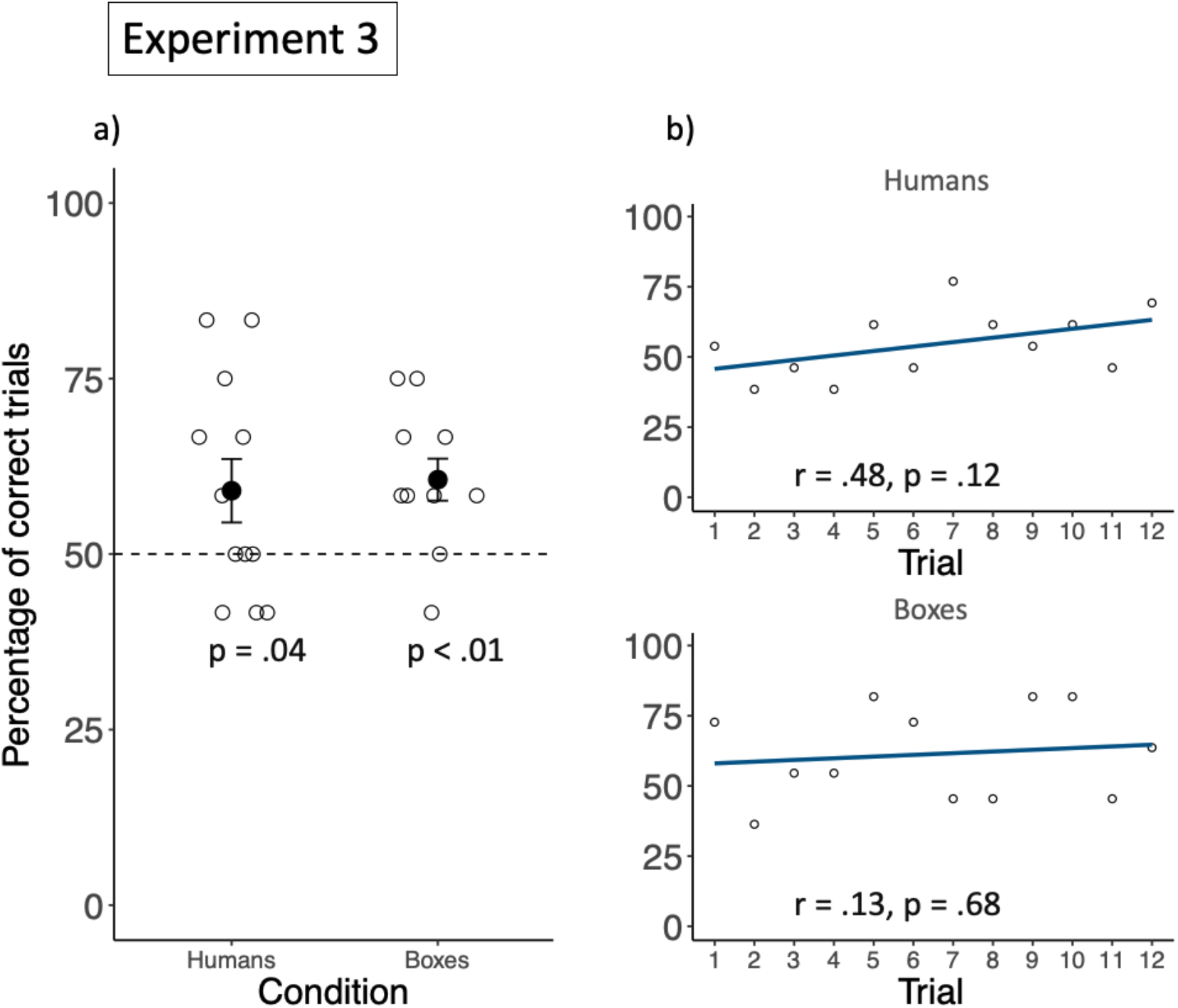
Group performance in Experiment 3. a) shows the group mean percentage of choosing the favourable option, in both conditions. “Humans” stands for the condition in which the rewarding patterns covaried with the social agents (Exp. 3a) and “Boxes” stands for the condition in which the rewarding patterns covaried with the boxes. Filled circles represent the group means, and the black back bars represent the standard errors. The empty circles represent individual performances. b) shows the correlation between group performance and trial number for each condition. The dark lines are regression lines and the light lines show the closest fit to the data points.

### Alternative strategy

An alternative to long-tailed macaques basing their decisions on the information provided in sampling phase is that they based their decisions on the outcomes of the previous trials and engaged in a Win-Stay, Loose-Switch strategy (hereafter WS-LS strategy). To disentangle between both possibilities, we ran for each condition of each experiment a post-hoc Generalised Linear Mixed Model (GLMM). We first coded, for each trial, whether individuals stayed with their choice in the next trial, or switched (Strategy variable). This resulted in 11 trials per individuals, as this information was not available for trial 12 (there was no trial 13). We also coded for each trial whether monkeys received a reward or not (Reward variable), and what the favourable choice would have been (Correct choice variable). For each model, we used as dependant variable Strategy, as fixed factors Reward and Correct choice and as random factor the individual IDs. If monkeys based their choices on previous outcomes, then Reward should be predictive of their stay/switch behaviour. If monkeys based their choices on the statistical information sampled in the sampling phase, then Correct choice should be predictive of their stay/switch behaviour. Results indicate that Correct choice was predictive of their stay/switch behaviour for each condition of each experiment, except Exp. 2b, in which Reward was predictive (see Table S4 from the supplementary material). These results suggest that in general, long-tailed macaques relied on the statistical information extracted from the sampling phase, but that in some specific cases, they might have switched to a WS-LS strategy.

### General discussion

Overall, our results suggest that long-tailed macaques extract statistical contingencies between patterns of rewards and different factors from repeated types of events, to assess which factors predict more probable rewards and adjust their decisions accordingly. Long-tailed macaques’ good performance in all conditions of Experiment 1 indicates that they did not rely on quantity heuristics to direct their choices, but rather used relative frequencies of rewards to estimate probabilities of future events. Our results also suggest that the condition – and therefore different contrasts of rewarding frequencies and different quantities of information – had an effect on monkeys’ probability to choose the favourable object. The monkeys’ confidence in choosing the favourable option seemed to increase with larger sample sizes.

The long-tailed macaques also seemed to rely on statistical information to make predictions in more complex scenarios involving several potential factors of different natures (social vs. non-social). However, the complexity or the nature of the factors seemed to affect monkeys’ ability to update their inferences with new information. In fact, the long-tailed macaques’ propensity to select the favourable option was lower in Experiments 2 and 3 than in Experiment 1, and did not increase across sessions. One reason for this could be that, with an increasing number of factors to consider, and with longer actions, monkeys had to test more hypotheses regarding which factors predicted more likely rewards, which in turn required more information and thus more sessions. Another reason could be that the nature of the potential factors prevented monkeys from extracting the relevant information, either because they do not possess the required mental representations, or because they did not realize that these factors were the same across sessions, and were therefore predictive within and between sessions. This might in turn have increased their propensity to rely on alternative strategies, such as a Win-Stay, Loose-Switch strategy based on previous outcomes. Our results suggest that this was the case for one of the conditions of Experiment 2, in which the factors predicting the rewards were two humans.

These findings suggest that not only apes, but also monkeys do engage in statistical reasoning. However, there seems to be differences between the reasoning ability required to solve tasks in which statistical information is extracted from repeated types of events, and tasks akin to games of chance. In fact, long-tailed macaques from the same population that we tested in the current study previously failed in tasks in which they had to make predictions from populations of objects to randomly drawn samples (see Placì et al. 2018). These differences could be due to the way statistical information had to be acquired in both types of tasks. In the current study, statistical information had to be extracted from the temporal contingencies between experienced events. In the previous study, assumptions about the sampling process had first to be extracted from visual cues provided by the experimenter (e.g., looking away from the buckets or closing her eyes and drawing an item out of each bucket). Once the sampling process was assumed to be random, the proportions of the different objects present in the buckets had to be estimated to compute the probability that an object of each type would be sampled. Monkeys did therefore not directly experience the probabilistic contingencies necessary to make rational decisions.

Interestingly, differences in statistical reasoning abilities due to different kinds of tasks have also been highlighted in human adults. Human adults are better at tasks in which they can extract statistical information from experienced events than in tasks in which they have to extract information from symbolic descriptions (see for example, Hertwig, Hogarth & Lejarraga, 2018; Rehder & Waldmann 2017). It seems that experiencing probabilistic contingencies improves decision making under uncertainty.

This difference in performance depending on the type of task could point to different reasoning mechanisms, one for solving probabilistic problems without experience and relying on information extracted in the present, and one for solving probabilistic problems with experience, relying on information extracted from a succession of events. Findings from a study done with 3- and 5-years-old children suggest that two cognitive mechanisms are indeed at play, as children’s reasoning about experienced events seemed to work independently from their reasoning about probabilities of single-events (Téglás et al., 2007). It would be interesting to investigate statistical reasoning with repeated events in more species, to investigate the evolutionary origins of this ability. Such ability has recently been investigated in a species of birds (Roberts et al 2018) with similar results to ours.

## Data availability statement

Supporting data will be available on request

## Acknowledgments

We thank Daniela Fuchs for her reliability assessment, Carolin Kade and Lukas Schad for their help with the data collection, Ludwig Ehrenreich for his technical assistance, all animal caretakers at the German Primate Center who assisted us and allowed us to conduct the experiments.

## References

1. Tenenbaum, J. B., Kemp, C., Griffiths, T. L., & Goodman, N. D. (2011). How to grow a mind: Statistics, structure, and abstraction. science, 331(6022), 1279–1285.

2. Jaynes, E. T. (2003). Probability theory: The logic of science. Cambridge university press.

3. Téglás E, Girotto V, Gonzalez M, & Bonatti LL. (2007). Intuitions of probabilities shape expectations about the future at 12 months and beyond. Proc. Natl. Acad. Sci. U. S. A. 104, 19156–9. (doi: 10.1073/pnas.0700271104)

4. Téglás E, Vul E, Girotto V, Gonzalez M, Tenenbaum JB, & Bonatti LL. (2011). Pure reasoning in 12-month-olds as probabilistic inference. Science (80-.). 332, 1054–1059. (doi: 10.1126/science.1196404)

5. Téglás, E., Ibanez-Lillo, A., Costa, A., & Bonatti, L. L. (2015). Numerical representations and intuitions of probabilities at 12 months. Developmental science, 18(2), 183–193.

6. Xu, F., & Garcia, V. (2008). Intuitive statistics by 8-month-old infants. Proceedings of the National Academy of Sciences, 105(13), 5012–5015

7. Xu, F., & Denison, S. (2009). Statistical inference and sensitivity to sampling in 11-month-old infants. Cognition, 112(1), 97–104

8. Denison S, & Xu F. (2014). The origins of probabilistic inference in human infants. Cognition 130, 335–347. (doi: 10.1016/j.cognition.2013.12.001)

9. Denison S, Trikutam P, & Xu F. (2014). Probability versus representativeness in infancy: can infants use naïve physics to adjust population base rates in probabilistic inference? Dev. Psychol. 50, 2009–2019. (doi: 10.1037/a0037158)

10. Gweon H, Tenenbaum JB, & Schulz LE. (2010). Infants consider both the sample and the sampling process in inductive generalization. Proc. Natl. Acad. Sci. 107, 9066–9071. (doi: 10.1073/pnas.1003095107)

11. Rakoczy H, Clüver A, Sauckle L, Stoffregen N, Gräbener A, Migura J, & Call J. 2014. Apes are intuitive statisticians. Cognition 131, 60–68. (doi: 10.1016/j.cognition.2013.12.011)

12. Eckert, J., Rakoczy, H., Call, J., Herrmann, E., & Hanus, D. (2018). Chimpanzees consider humans’ psychological states when drawing statistical inferences. Current Biology

13. Eckert, J., Call, J., Hermes, J., Herrmann, E., & Rakoczy, H. (2018). Intuitive statistical inferences in chimpanzees and humans follow Weber’s law. Cognition, 180, 99–107

14. Tecwyn EC, Denison S, Messer EJE, & Buchsbaum D. (2017). Intuitive probabilistic inference in capuchin monkeys. Anim. Cogn. 20, 243–256. (doi: 10.1007/s10071-016-1043-9)

15. Placì, S., Eckert, J., Rakoczy, H., & Fischer, J. (2018). Long-tailed macaques (Macaca fascicularis) can use simple heuristics but fail at drawing statistical inferences from populations to samples. Open Science, 5(9), 181025

16. Eckert, J., Rakoczy, H., & Call, J. (2017). Are great apes able to reason from multi-item samples to populations of food items?. American journal of primatology, 79(10), e22693

17. Waismeyer, A., Meltzoff, A. N., & Gopnik, A. (2015). Causal learning from probabilistic events in 24-month-olds: an action measure. Developmental science, 18(1), 175–182

18. Gweon, H., & Schulz, L. (2011). 16-month-olds rationally infer causes of failed actions. Science, 332(6037), 1524–1524

19. Gopnik, A., Sobel, D. M., Schulz, L. E., & Glymour, C. (2001). Causal learning mechanisms in very young children: two-, three-, and four-year-olds infer causal relations from patterns of variation and covariation. Developmental psychology, 37(5), 620

20. Völter, C. J., Sentís, I., & Call, J. (2016). Great apes and children infer causal relations from patterns of variation and covariation. Cognition, 155, 30–43

21. Edwards, B. J., Rottman, B. M., Shankar, M., Betzler, R., Chituc, V., Rodriguez, R & Santos, L. R. (2014). Do capuchin monkeys (Cebus apella) diagnose causal relations in the absence of a direct reward?. PloS one, 9(2), e88595.

22. Baayen, R. H. (2010). Analyzing linguistic data: A practical introduction to statistics. Cambridge: Cambridge University Press.

23. Rehder, B., & Waldmann, M. R. (2017). Failures of explaining away and screening off in described versus experienced causal learning scenarios. Memory & cognition, 45(2), 245–260.

24. Hertwig, R., Hogarth, R. M., & Lejarraga, T. (2018). Experience and description: Exploring two paths to knowledge. Current Directions in Psychological Science, 27(2), 123–128

25. Roberts, W. A., MacDonald, H., & Lo, K. H. (2018). Pigeons play the percentages: computation of probability in a bird. Animal Cognition, 1–7

