## Supplementary material for "Long-tailed macaques extract statistical information from repeated types of events to make rational decisions under uncertainty"

*Corresponding Author: Sarah Placì

[+]Equal contribution

Supplementary material

Table S1. Individual participation and performance in each experiment. For each condition of each experiment, we report the number of times, on a total of twelve test trial, that each individual selected the favourable option. Bars indicate conditions in which individuals did not participate.

| | **Experiment 1** | | | | **Experiment 2** | | **Experiment 3** | |
|---|---|---|---|---|---|---|---|---|
| | 10/10 vs. 4/10 | 8/10 vs. 4/10 | 4/4 vs. 4/10 | 4/4 vs. 4/10 (side control) | 6/6 vs. 6/12 | 4/4 vs. 4/10 | Humans | Boxes |
| Ilia | 10* | 10* | 12* | 11* | 8 | 6 | 5 | 7 |
| Linus | 12* | 7 | 7 | 7 | 6 | 5 | 5 | 9 |
| Lukas | 10* | 8 | 4 | 8 | 7 | 6 | 8 | 8 |
| Mara | 11* | 10* | 7 | 5 | 9 | 5 | 6 | 7 |
| Mars | 11* | 9* | 11* | 8 | 5 | / | 6 | 9 |
| Max | 11* | 12* | 11* | 10* | 9 | 9 | 10* | 7 |
| Maja | 10* | / | / | 6 | 7 | 8 | 6 | 7 |
| Meiwi | 10* | 8 | 5 | 8 | 8 | 5 | 10* | 5 |
| Mila | 11* | 10* | 7 | 9 | 8 | 8 | 9 | 7 |
| Milka | 10* | 11* | 8 | 8 | 6 | 8 | 5 | 8 |
| Moritz | / | / | / | 9 | / | 6 | 7 | / |
| Paul | 12* | / | / | / | / | / | / | / |
| Sambia | / | / | / | 9 | 10* | 6 | 8 | 6 |
| Snickers | 10* | 11* | 12* | / | 9 | 11* | / | / |
| Sissi | / | / | / | / | / | 5 | / | / |
| Smilla | / | / | / | 10* | / | 9 | / | / |

Experiment 1 - Generalized linear mixed model

We ran a GLMM to assess whether the rewarding frequencies had an effect on monkeys´ performance. As Exp. 1d served as a control to rule out that monkeys only associated the rewarding frequencies with the side of presentation, but that otherwise it had the same rewarding frequencies as Exp. 1c, we did not include it in this analyse. As dependant variable, we used monkeys´ choices (1 for choosing the favourable option, and 0 for choosing the unfavourable one). As fixed factors, we used the condition (Exp. 1a, Exp. 1b and Exp. 1c), the trial number, that we beforehand z-transformed, and the order in which each individual underwent the different condition, which was also z-transformed. As random variable, we used the individual IDs.

The model was the following: Choice ~ Condition + z.Trial + z.Order + (1I Individual)

We used the “vif” function of the “car” package to check for collinearity. There was no collinearity between our predictors, as all values were below 3. Our model was not overdispersed, as the dispersion parameter was 0.856. We used the “drop1” function of the package “lme4” using “Chisq” to calculate the p-value. As we were also interested in testing differences between conditions, we ran a postdoc analyse using the function “glht from the package “multicomp”. Results revealed that condition had a main effect on subjects´ performance, as well as the trial number (see Table S2), and that there was a significant difference between Exp. 1a and Exp. 1c (see Table S3).

Table S2. Results of the generalized linear mixed model.

| | Estimate | Std. Error | Chisq | p-value |
|---|---|---|---|---|
| Intercept | 3.374 | 0.551 | * | * |
| ConditionExp1b | -1.328 | 0.661 | 14.457 | 0.001 |
| ConditionExp1c | -2.311 | 0.689 | | |
| z.Trial | 1.640 | 0.214 | 94.615 | 0.000 |
| z.Order | 0.137 | 0.300 | 0.207 | 0.649 |

Table S3. Results of the multicomparison analyse between conditions.

| | Estimate | Std. Error | z value | p-value |
|---|---|---|---|---|
| ConditionExp1b == 0 | -1.328 | 0.661 | -2.009 | 0.10 |
| ConditionExp1c == 0 | -2.311 | 0.689 | -3.354 | 0.002 |
| ConditionExp1b – ConditionExp1c == 0 | 0.983 | 0.372 | 2.64 | 0.021 |

Table S4. This table shows the results of the generalised linear mixed models, in which the stay/switch behaviour of the monkeys was the dependant variable, and whether they got a reward or not (Reward) and whether they chose the favourable option (Correct_Choice) were the fixed factors and individuals IDs were included as random factors.

| | | Estimate | Std. Error | z value | p-value |
|---|---|---|---|---|---|
| Exp. 1a | Intercept | -2.16 | 1.15 | -1.87 | 0.06 |
| | Reward | 0.11 | 1.62 | 0.07 | 0.94 |
| | Correct_Choice | 4.79 | 1.38 | 3.46 | <.001 |
| Exp. 1b | Intercept | -0.24 | 0.50 | -0.48 | 0.63 |
| | Reward | 0.16 | 0.61 | 0.26 | 0.80 |
| | Correct_Choice | 2.54 | 0.60 | 4.26 | <.001 |
| Exp. 1c | Intercept | 0.90 | 0.57 | 1.58 | 0.12 |
| | Reward | -0.22 | 0.79 | -0.27 | 0.78 |
| | Correct_Choice | 2.18 | 0.80 | 2.72 | <.01 |
| Exp. 1d | Intercept | -0.69 | 0.37 | -1.88 | 0.06 |
| | Reward | 0.11 | 0.67 | 0.16 | 0.88 |
| | Correct_Choice | 1.63 | 0.60 | 2.70 | <.01 |
| Exp. 2a | Intercept | -0.70 | 0.45 | -1.56 | 0.12 |
| | Reward | 0.02 | 0.64 | 0.03 | 0.97 |
| | Correct_Choice | 1.02 | 0.52 | 1.96 | 0.05 |
| Exp. 2b | Intercept | -0.80 | 0.33 | -2.40 | 0.02 |
| | Reward | 1.24 | 0.54 | 2.30 | 0.02 |
| | Correct_Choice | -0.15 | 0.48 | -0.31 | 0.76 |
| Exp. 3a | Intercept | -0.27 | 0.34 | -0.78 | 0.43 |
| | Reward | -0.50 | 0.60 | -0.83 | 0.41 |
| | Correct_Choice | 1.09 | 0.55 | 1.96 | 0.05 |
| Exp. 3b | Intercept | -0.19 | 0.36 | -0.54 | 0.59 |
| | Reward | -1.27 | 0.74 | -1.73 | 0.08 |
| | Correct_Choice | 1.74 | 0.68 | 2.55 | 0.01 |

Alternative strategy

We ran for each condition of each experiment a post-hoc Generalised Linear Mixed Model (GLMM). We first coded, for each trial, whether individuals stayed with their choice in the next trial, or switched (Strategy variable). This resulted in 11 trials per individuals, as this information was not available for trial 12 (there was no trial 13). We also coded for each trial whether monkeys received a reward or not (Reward variable), and what the favourable choice would have been (Correct choice variable). For each model, we used as dependant variable Strategy, as fixed factors Reward and Correct choice and as random factor the individual IDs.

The model was the following: Strategy ~ Reward + Correct_Choice + (1I Individual).

For each model, we used the “vif” function of the “car” package to check for collinearity. There was no collinearity between our predictors, as all values were below 3. Our models were not overdispersed, as the dispersion parameter were 0.76 for Exp. 1a, 1.04 for Exp. 1b, 0.83 for Exp. 1c, 1.03 for Exp. 1d, 1.00 for Exp. 2a, 1.03 for Exp. 2b, 1.00 for Exp. 3a, and 1.03 for Exp. 3b.
